# Extracellular RNA drives Electromethanogenesis in a Methanogenic Archaeon

**DOI:** 10.1101/2025.07.06.663362

**Authors:** Rhitu Kotoky, Obinna Markraphael Ajunwa, Satoshi Kawaichi, Haluk Beyenal, Jerome T. Babauta, Rikke Louise Meyer, Amelia-Elena Rotaru

**Affiliations:** Nordcee, Department of Biology, University of Southern Denmark, Odense, Denmark; Interdisciplinary Nanoscience Center (iNANO), Aarhus University, Aarhus, Denmark; Center for Electromicrobiology, Department of Biology, Aarhus University, Aarhus, Denmark; Department of Biology, Aarhus University, Aarhus, Denmark; The School of Chemical engineering and Bioengineering, Voiland College of Engineering and Architecture, Washington State University, Pullman, Washington, USA; Gamry Instruments Inc, Warminster, PA, USA

## Abstract

Methanogenic archaea account for two-thirds of global methane emissions. Some species, including *Methanosarcina barkeri*, reduce CO_2_ by directly acquiring electrons from solid substrates. However, the mechanism of electron acquisition in *M. barkeri* has remained unclear because this archaeon lacks the multiheme c-type cytochromes that drive extracellular electron transfer in many other microbes. Here we show that *M. barkeri* releases abundant extracellular nucleic acids during early growth, primarily short RNAs (78%). These extracellular nucleic acids assemble into G-quadruplexes (G4s) and B-DNA architectures that decorate cell surfaces and link aggregates. Surface-associated G4s are folded *in vivo* in a conformation compatible with cofactor binding and redox chemistry. Enzymatic degradation of extracellular nucleic acids abolished electron uptake and electromethanogenesis, whereas addition of synthetic G4-RNAs doubled methane yields and lowered cell-electrode interfacial resistance. These effects were not observed when cells were grown on soluble substrates. Together, these findings identify eRNA as a previously unrecognized electron conduit in methanogens, raising the possibility that RNA-based electron transfer may predate more elaborate protein-based electron conduits, with implications for models of early earth metabolism and for the design of next-generation bioenergy systems.

## Main

Methanogenic archaea are an ancient lineage that thrives in anoxic environments and accounts for most biogenic methane production on Earth. In these settings, methanogens can acquire electrons not only from syntrophic partner cells^1,2^ but also from conductive materials, including minerals,^3,4^ metals^5–7^ and electrodes^8,9^. This capacity, mediated either through diffusible carriers or by direct extracellular electron transfer (EET), enables methane production when soluble electron donors are limiting or absent. Yet, how methanogenic archaea perform EET remains unknown in most species.

In electroactive bacteria, EET is typically mediated by cell surface multiheme c-type cytochromes (MHCs), either as discrete outer-surface components, as polymerized conductive ‘nanowires’, or in association with type IV pili^10–12^. An MHC-dependent EET mechanism^13,14^ has also been demonstrated in the methanogen *Methanosarcina acetivorans*^15,16^. However, most methanogenic archaea lack these canonical surface electron-transfer components, indicating that they must rely on alternative strategies for EET.

This gap is particularly evident in species of the order *Methanosarcinales*, which includes *Methanosarcina* and *Methanothrix*, that can engage in direct interspecies electron transfer (DIET) with electrogenic bacteria, or can acquire electrons from minerals, metals, or electrodes^1,17,18^. A subset of *Methanosarcina* (Type II) contain the multiheme cytochrome MmcA and rely on it for EET^19,20^. In contrast, Type I *Methanosarcina* (without MmcA), and all *Methanothrix* species are devoid of multiheme cytochromes altogether^21–23^ indicating their use of unknown alternative EET strategies.

One viable alternative is extracellular nucleic acids (eNAs), which can support electron transport through π-stacked bases in structures such as G-quadruplexes (G4)^24^. Recent studies in bacterial biofilms implicated eNAs in extracellular electron transfer in conjunction with defined redox-active molecules, including phenazines and hemin^25,26^. Yet whether eNAs can mediate direct extracellular electron uptake coupled to energy metabolism has remained unknown. This question is especially relevant in methanogens, in which extracellular electron uptake drives methane formation ^1,2,8,9,17,18,27^.

Here we show that *Methanosarcina barkeri* strain MS (DSM 800), a model methanogen lacking canonical MHC-based EET machinery, relies on self-generated, surface-associated extracellular RNA (eRNA) organized in G4 superstructures for cathodic electron uptake coupled to CO_2_ reduction to methane. These findings establish extracellular RNA as a previously unrecognized basis for methanogenic electron uptake and uncover an unrecognized role for extracellular RNA in archaeal energy metabolism.

### Extracellular nucleic acids self-organize into G4 structures that support redox chemistry at the cell surface

To determine whether *M. barkeri* releases extracellular nucleic acids (eNAs) during methanogenic growth, we quantified eNAs alongside biomass accumulation and methane production. Total eNAs (<50 bp; Extended Data Fig. 1) accumulated sharply at growth onset, coincident with the initial rise in biomass and methane production (Fig. 1a). This early surge was detected by both spectrophotometric and fluorometric measurements, reaching 35 ± 1.6 and 11 ± 0.3 µg mL^-1^ culture, respectively. Fluorometric analysis further showed that the extracellular nucleic acid pool was dominated by eRNA, which comprised 78% of total eNAs, whereas single-and double-stranded DNA together accounted for <20% (Fig. 1b, inset). These results identify a predominantly RNA extracellular nucleic acid pool released at growth onset and prompted us to test whether eNAs localize at the cell surface.

**Fig. 1.**
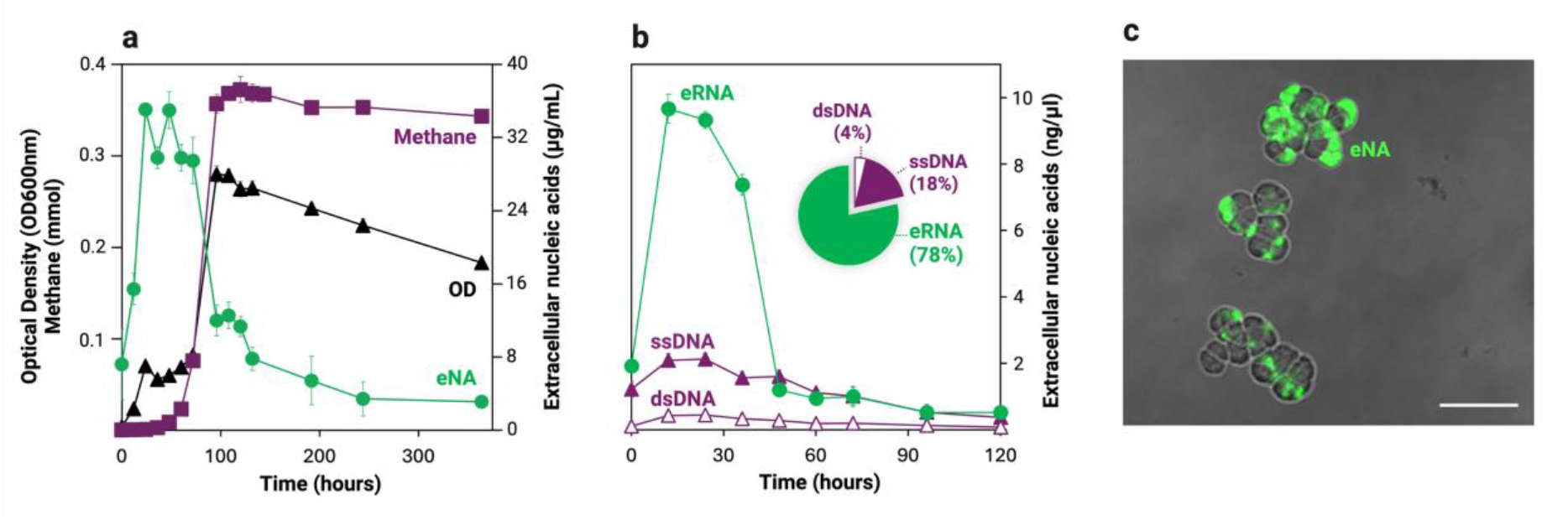
Extracellular nucleic acids (eNAs) accumulate during early growth of *M. barkeri*. **a**, Accumulation of total eNAs in the culture medium alongside optical density at 600 nm and methane production. **b**, Fluorometric speciation of extracellular RNA (eRNA, green), single-stranded DNA (ssDNA, solid purple), and double-stranded DNA (dsDNA, open purple) over time. Inset, relative composition of eNAs at peak abundance (12-36 h). Data are mean ± SD for n≥3 independent cultures. **c**, Confocal micrograph of live *M. barkeri* cells stained with a membrane-impermeable nucleic acid dye TOTO-1 (green), showing eNAs surrounding cells. Scale bar, 10 µm. The confocal micrograph is representative of three independent cultures. Created in BioRender. Rotaru, A. (2026) https://BioRender.com/vvxhzbz

Confocal imaging of intact *M. barkeri* cells stained with cell-impermeant dyes revealed a pericellular fluorescent eNA halo surrounding cells (Fig. 1c), while dual staining of membranes and eNAs confirmed membrane integrity as the nucleic acid signal remained external to the membrane (Extended Data Fig. 2a-c). Together, these observations show that the released eNAs are retained outside intact cells and remain closely associated with the cell surface.

We next asked whether *M. barkeri’s* eNAs adopt the B-DNA and G-quadruplexes (G4s) superstructures implicated in EET in bacterial biofilms^24,25,28^. Genome analyses identified 1,685 predicted G4-forming sequence signatures in the *M. barkeri* genome, supporting its capacity to form G4-rich structures^29^. Structure-specific immunolabeling^26^ detected both B-DNA and G4s surrounding cells within aggregates and in structures linking neighboring aggregates (Fig. 2a). Notably, G4-rich foci were often associated with long B-DNA filaments (>10 µm) in a bead-on-a-string arrangement (Fig. 2), resembling he EET machinery of electroactive bacteria, namely pili decorated with multiheme cytochromes^30^.

**Fig. 2.**
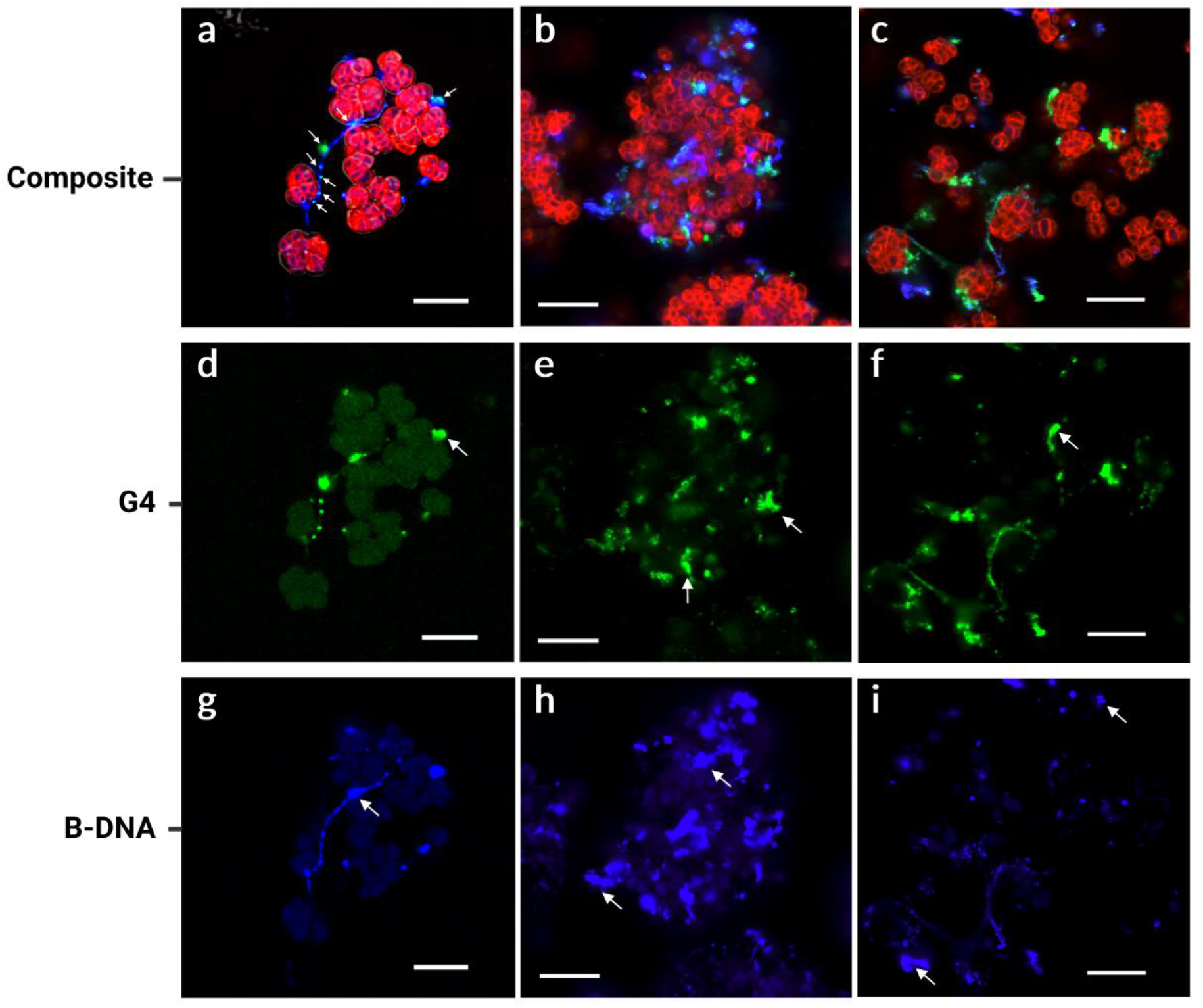
Extracellular B-DNA and G-quadruplex structures associated with *M. barkeri* aggregates. **a-c**, Confocal micrographs of intact *M. barkeri* cells triple-labeled to visualize cell membranes (FM4-64, red), B-DNA (AlexaFluor-405, blue) and G-quadruplexes (Atto-488, green). In **a**, white arrows indicate G4-rich foci associated with a B-DNA filament. **d-f**, Single-channel micrographs of immunolabeled G4 structures. **g-i**, Single-channel micrographs of immunolabeled B-DNA structures. Scale bars, 10 μm. Confocal images are representative of three independent cultures. Created in BioRender. Rotaru, A. (2026) https://BioRender.com/fjiebsf

Because G4 structures must be correctly folded to support cofactor binding and redox chemistry, we next used a hemin-dependent Tyramide Signal Amplification (TSA) assay to test whether surface-associated G4s of *M. barkeri* are natively folded *in vivo*. Correctly folded G4s bind hemin to form DNAzyme/RNAzyme complexes that catalyze H_2_O_2_-dependent oxidation of tyramide to radicals that deposit locally yielding a fluorescent signal^26^. Hemin-supplemented *M. barkeri* cells exhibited strong fluorescence at the cell surface, indicating that surface-associated G4s were correctly folded to bind hemin and form catalytically active complexes. The signal formed a diffuse sheath outlining most cells within aggregates (Fig. 3a-c, Extended Data Fig. 3). Together, these observations show that surface-associated G4s in *M. barkeri* are natively folded in a manner compatible with cofactor binding and redox chemistry at the cell surface.

**Fig. 3.**
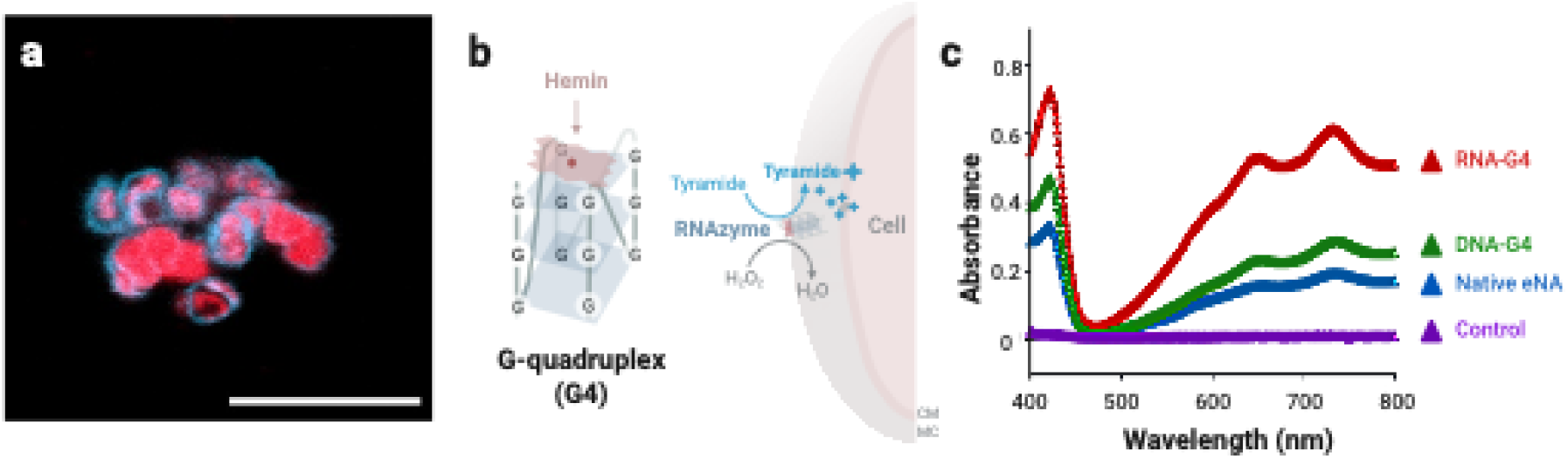
Hemin-dependent DNAzyme/RNAzyme activity of surface associated G4s in *M. barkeri*. **a**, Confocal micrographs of a hemin-dependent *in vivo* tyramide signal amplification (TSA) assay on *M. barkeri* cells. Deposited tyramides (cyan) mark extracellular G4-hemin complexes with DNAzyme/RNAzyme activity surrounding cells counterstained with the intracellular nucleic acid stain SYTO60 (red). Scale bars, 10μm. Additional controls are shown in Extended Data Fig. 3. **b**, Schematic of a G4-hemin complex with DNAzyme/RNAzyme activity catalyzing H_2_O_2_-dependent oxidation of tyramide to reactive radicals that covalently deposit on the tyrosines of nearby proteins at the cell surface. **c**, *In-vitro* peroxidase assays showing absorbance spectra (400-800 nm) of oxidized ABTS generated by pre-folded nucleic acids: synthetic G4-RNA (red), synthetic G4-DNA (green), native eNAs from *M. barkeri* (blue), and a negative control lacking G4s (purple). MC: methanochondroitin, CM: cell membrane. Created in BioRender. Rotaru, A. (2026) https://BioRender.com/w1fl488

### Extracellular nucleic acids are required for electromethanogenesis in *M. barkeri*

*M. barkeri* performs electromethanogenesis when provided with a cathode as the sole electron donor ^8,9,31^. Under cathodic growth conditions at -430 mV versus the standard hydrogen electrode (SHE), *M. barkeri* aggregates attached to carbon felt electrodes were embedded within extracellular G4 and B-DNA structures (Fig. 4a). Substantial G4 and B-DNA deposits persist on the carbon felt surface even when aggregates detach, showing that eNAs remain firmly associated with the electrode surface during electromethanogenesis. Consistent with active electromethanogenesis, *M. barkeri-*inoculated reactors generated a sustained cathodic current relative to abiotic control (Fig. 4b), accompanied by methane accumulation in the reactor headspace.

**Fig. 4.**
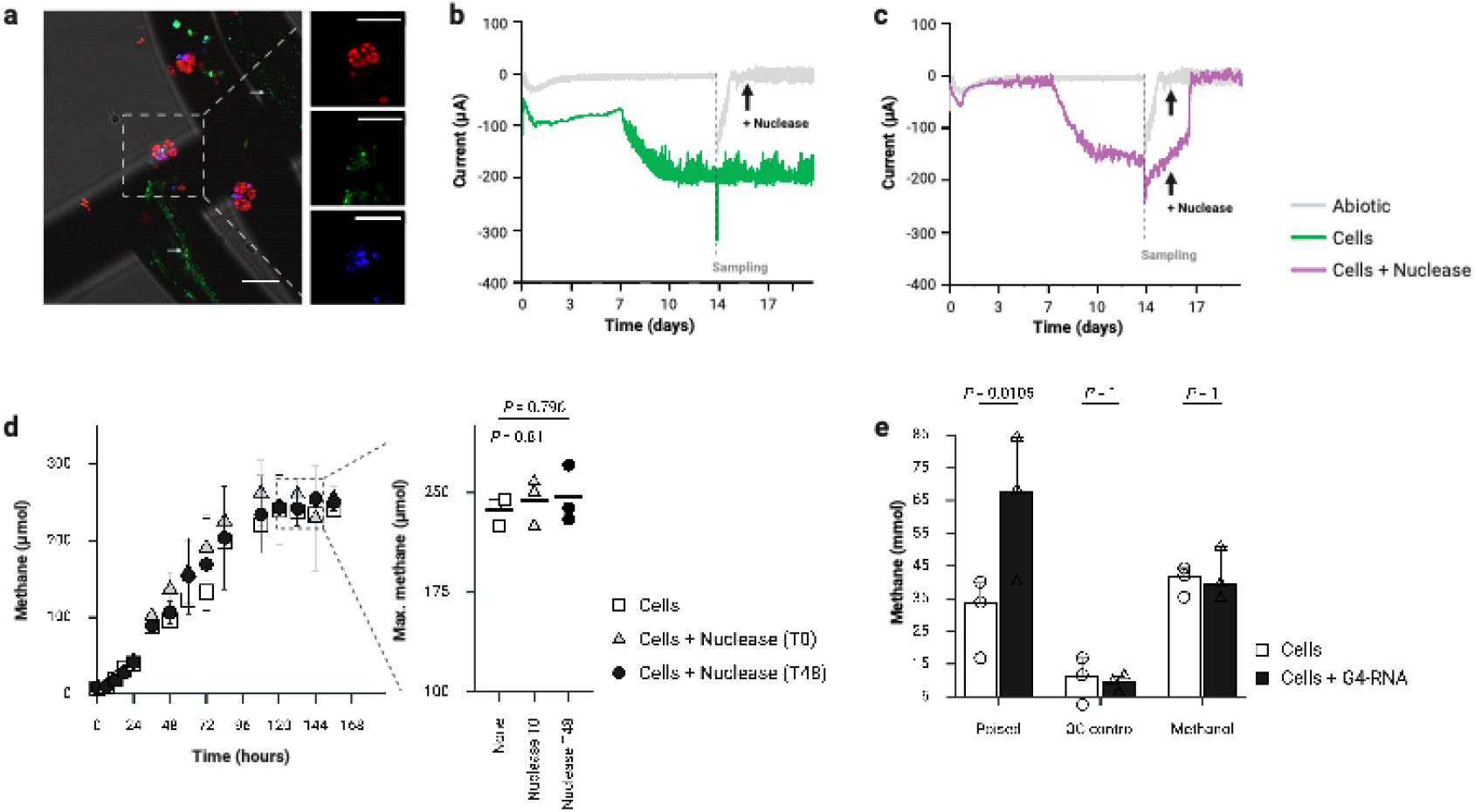
Effects of extracellular nucleic acid degradation and synthetic G4-RNA amendments on electromethanogenesis by *M. barkeri*. **a**, Confocal micrographs of *M. barkeri* grown on carbon felt electrodes poised at -430 mV versus SHE, labelled for membranes (FM4-64, red), B-DNA (AlexaFluor-405, blue), and G4 structures (Atto-488, green). Insets show the individual fluorescence channels. White arrows indicate G4-rich deposits on the electrode surface. Scale bars, 10 μm. **b**, Chronoamperometric trace showing cathodic current produced by *M. barkeri* cells (green), abiotic controls (gray) showing baseline current. **c**, Chronoamperometric trace showing rapid loss of cathodic current after addition of micrococcal nuclease on day 16 (purple). Abiotic controls (gray) remained unaffected. Arrows indicate nuclease addition; dashed lines indicate gas sampling. **d**, Methane production in methanol-fed cultures with no nuclease or with nuclease added at inoculation (T0) or after 48 h (T48). Inset, maximum methane yields across treatments between 120-144 h. **e**, Methane production by *M. barkeri* under poised-cathode, open-circuit and methanol-grown conditions in the presence or absence of synthetic G4-RNAs. Data are mean ± SD for biological triplicates (n≥3). Images and chronoamperometric traces are representative of n≥3 biological replicates. Created in BioRender. Rotaru, A. (2026) https://BioRender.com/qvsc7lv structures such as G-quadruplexes^32^. Within hours of adding the micrococcal nuclease, the cathodic current fell to baseline levels and no further methane production was detected (Fig. 4c, Extended Data Fig. 6). Crucially, the same nuclease addition did not inhibit growth of methanol-fed cultures (Fig. 4d), demonstrating that eNA degradation selectively disrupts electron uptake from a cathode, but does not disrupt growth on soluble substrates.

To determine whether eNAs are necessary for electromethanogenesis, we depleted extracellular nucleic acids when *M. barkeri* was at peak current density and methane production. Micrococcal nuclease digests both extracellular DNA and RNA indiscriminately, including their secondary An alternative explanation for the collapse in cathodic electron uptake was that micrococcal nuclease impaired cell attachment to the cathode. To test this, we quantified biomass retention on carbon felt electrodes in the presence and absence of micrococcal nuclease. Total electrode-associated protein did not differ after nuclease treatment (Extended Data Fig. 7), indicating that the loss of cathodic electron uptake was not caused by reduced biomass retention. These results show that depletion of eNAs disrupts electromethanogenesis without impairing cell attachment to electrodes, supporting a direct requirement for eNAs in cathodic electron uptake.

Although eRNA dominates the extracellular nucleic acid pool, micrococcal nuclease degrades extracellular DNA and RNA indiscriminately, leaving the specific contribution of G4-RNAs to electromethanogenesis unresolved. We therefore tested whether synthetic G4-RNAs enhance electromethanogenesis by *M. barkeri* on cathodes poised at -430 mV vs SHE, alongside open circuit and methanol-grown controls. Remarkably, under poised conditions, synthetic G4-RNAs doubled cathodic methane production (Fig. 4e, n=3, p=0.01). This effect was absent in open circuit and methanol-grown conditions (Extended Data Fig. 5, Fig. 4e), ruling out nonspecific metabolic stimulation. These results indicate that exogenous G4-RNAs specifically enhance cathodic electron uptake rather than altering core methanogenic metabolism.

Electrochemical impedance spectroscopy (EIS) was used to test whether G4-RNA alters electron transfer at the cell-electrode interface^33,34^. Nyquist plots of carbon printed electrodes bearing *M. barkeri* cells showed broad, depressed arcs that became smaller in the presence of exogenous G4-RNA, whereas these features were absent in abiotic controls (Fig. 5a). At low frequencies, the arcs transition to a tail-like feature consistent with mass-transport or pseudocapacitive contributions, which were accounted for in the equivalent-circuit model (Fig. 5c)^35^. Fitting to an equivalent circuit model showed that G4-RNA reduced the charge-transfer resistance at the cell-electrode interface from 9.9 kΩ to 4.2 kΩ (p = 0.001, n=4, Fig. 5c), whereas solution resistance and pseudo-biofilm resistance remained unchanged (Fig. 5c). Cyclic voltammetry provided complementary evidence of biologically associated cathodic activity on printed electrodes (Fig. 5b). Together, these data indicate that G4-RNA lowers the apparent interfacial charge-transfer barrier to electron transfer.

**Fig. 5.**
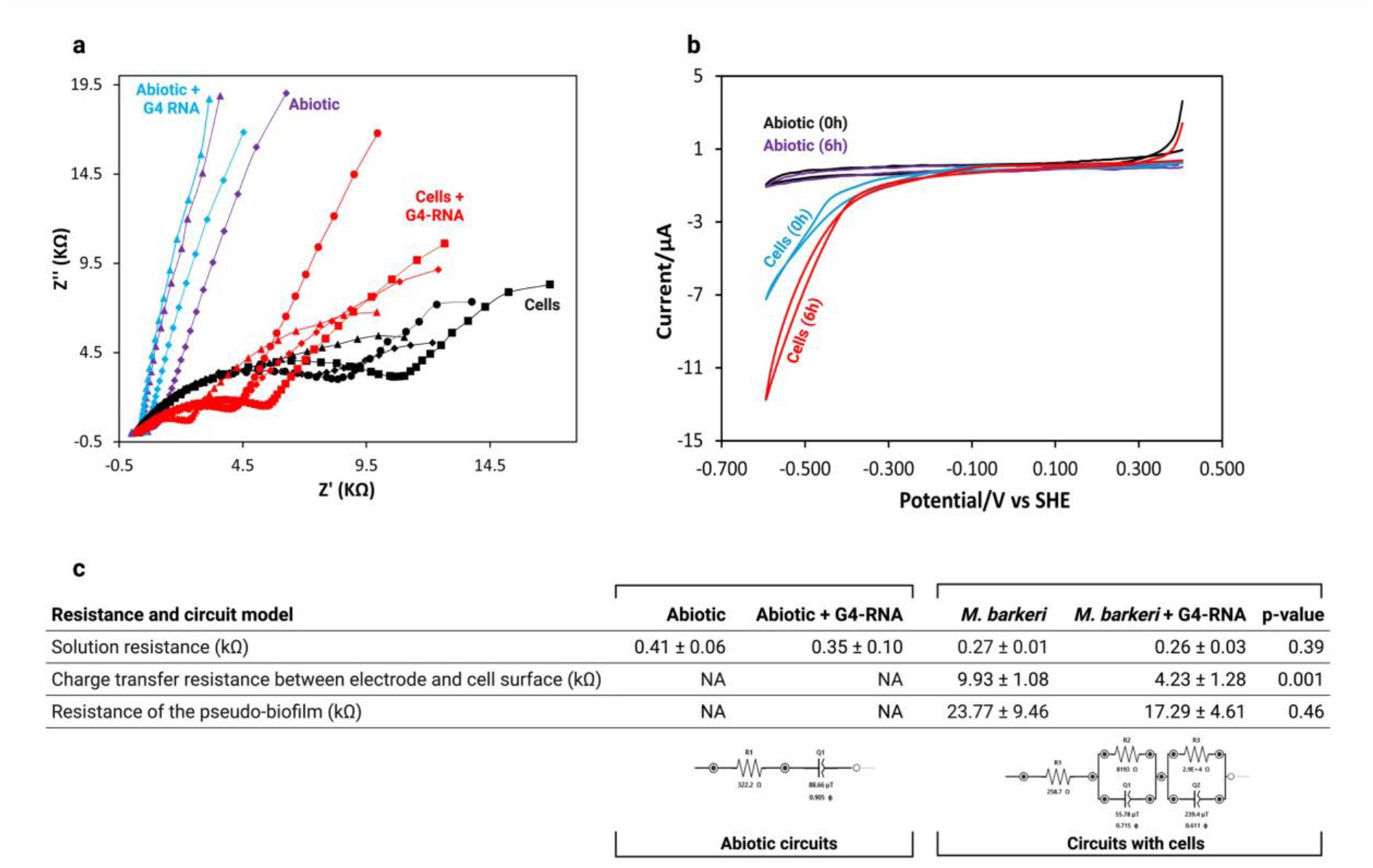
Electrochemical impedance spectroscopy (EIS) and cyclic voltammetry (CV) of *M. barkeri* on carbon printed electrodes. **a**, Nyquist plots showing the impedance response of abiotic electrodes, abiotic electrodes with added G4-RNA, electrodes bearing *M. barkeri* cells, and electrodes bearing *M. barkeri* cells with added G4-RNA. **b**, Cyclic voltammograms of carbon printed electrodes bearing *M. barkeri* cells recorded straightaway (0 h) and after 6 h of chronoamperometric operation, compared with abiotic controls. CVs are representative scans. **c**, Equivalent circuit models and fitted resistance values for solution resistance, charge transfer resistance at the cell-electrode interface, and pseudo-biofilm resistance. Values are mean ± SD of n = 4 biological replicates. Statistical significance was determined using a two-tailed t-test. The additional parameters and values from the fitted models have been given in Extended Data Table 1. Created in BioRender. Rotaru, A. (2026) https://BioRender.com/a1vf9pe

### Ecological and evolutionary implications of RNA-based electron uptake

Here, we identify an eRNA-based mode of electron uptake during electromethanogenesis in *M. barkeri*. Unlike bacteria, in which extracellular nucleic acids (primarily eDNA) accumulate late in growth or during biofilm dissolution^36–39^, *M. barkeri* releases substantial amounts of eRNA at growth onset. These extracellular nucleic acids assembled into superstructures around cell aggregates, with G4-rich elements coating cell surfaces in conformations compatible with redox chemistry. Conclusively, nuclease-mediated depletion of extracellular nucleic acids abolished electron uptake, whereas addition of G4-RNA enhanced electromethanogenesis and lowered cell-electrode interfacial resistance. Together, these findings establish G4-RNAs as electron-uptake conduits and expand the known repertoire of electron-transfer conduits beyond cytochrome nanowires and pilins.

Why does a relatively slow-growing anaerobe such as *M. barkeri* deploys eRNA, rather than eDNA, as an electrical conduit for EET remains unclear. One plausible explanation lies in the structural properties of G4-RNA. Compared to G4-DNA, G4-RNA adopts more rigid and thermodynamically stable conformations that favor intermolecular assembly and strong interactions with planar redox-active cofactors^40,41^. This greater structural stability may enhance binding to flat redox-active cofactors such as hemin, flavins and phenazine^42–44^. Consistent with this idea, our *in vitro* assays showed that G4-RNA displays greater peroxidase activity than sequence-identical G4-DNA (Fig. 3d), supporting a functional advantage of RNA-based architectures for redox chemistry.

A second, non-exclusive possibility is that eRNA reduces the biosynthetic burden of constructing a diverse extracellular electron transfer network from cell-surface proteins, including multiple classes of multiheme cytochromes. If so, an eRNA-dependent mechanism of EET may represent a simpler solution that preceded the evolution of more elaborate protein-based extracellular electron-transfer systems.

Understanding this mechanism is particularly relevant because *M. barkeri* and many Type I *Methanosarcina* lack MHCs and are broadly distributed across methane-producing environments, including wastewaters, anaerobic digesters, rice paddies and other anoxic aquatic habitats. In such settings, eRNA may influence how these methanogens interact with insoluble electron-donating surfaces, including conductive particles, other cells and electrodes. More broadly, these findings raise the possibility that eRNA-based electron uptake contributes to methane production in natural and engineered environments and may provide a new target for improving electromethanogenesis in bioelectrochemical systems that couple electrical current uptake to microbial CO_2_ reduction to methane.

## Methods

### Microorganisms, media, and cultivation conditions

*Methanosarcina barkeri* MS (DSM 800) was revived from the laboratory freezer stocks and cultivated anaerobically under an N_2_: CO_2_ (80:20, v/v) atmosphere in serum bottles containing DSM-120 medium sealed with butyl rubber stoppers. Cultures were incubated statically at 37°C. Routine growth was supported by 50 mM methanol as the electron donor. The reactor precultures were prepared with 10 mM acetate and 20 mM methanol.

For bio-electrochemical experiments, a modified DMS-120 reactor medium was prepared without any potential soluble electron shuttles, excluding resazurin, sodium sulfide, yeast extract, and tryptone. Under these conditions, the cathode served as the electron donor (see below).

### Analytical measurements

Methane (CH_4_) concentrations in the culture headspaces were measured using a Trace 1300 gas chromatograph (Thermo-Scientific) equipped with a TracePLOT™ TG-BOND MSieve 5A column and a Flame Ionization Detector (FID). Nitrogen was used as carrier gas at a flow rate of 1 mL min^-1^, and the injector, oven, and detectors were kept at 220°C, 40°C, and 230°C, respectively.

For electrochemical reactors, when both CH_4_ and H_2_ had to be quantified, we used the same GC and column fitted to a Thermal Conductivity Detector (TCD). In this case, argon served as carrier gas at a flow rate of 25 mL/min, with the injector, oven, and detector temperatures set at 150°C, 70°C, and 200°C respectively. Gas concentrations were converted to molar quantities using the ideal gas law under standard conditions.

### Bio-electrochemical reactor setup and perturbation experiments

The bio-electrochemical incubations were performed in dual-chambered H-cell reactors (Adams and Chittenden, USA), each chamber holding 100 mL medium and separated by a Nafion™ N117 proton exchange membrane (Ion Power). Carbon felt electrodes (2.5 × 2.5 × 1.2 cm, Thermo Scientific) served as the working electrode (cathode) and counter electrode (anode) and were connected via titanium wires. A leak-free Ag/AgCl reference electrode (3.4 M KCl) (CMA Microdialysis, Sweden) was positioned approximately 1 cm from the cathode.

The cathodes were poised at -430 mV versus the Standard Hydrogen Electrode (SHE) using a multichannel potentiostat (MultiEmStat Potentiostat, PalmSens, The Netherlands). Exponentially growing *M. barkeri* (OD600~0.2) were used to inoculate the cathodic chamber to a final inoculum of 20% (v/v). Open circuit controls were assembled identically but were not connected to the potentiostat.

To assess effects on cathodic electron uptake, we applied two perturbations: (i) amended the inoculum prior to reactor inoculation with additional synthetic G4-RNA (2.5 µM); and (ii) degraded the native G-quadruplexes with micrococcal nuclease (15 U mL^-1^) during peak current draw. This micrococcal nuclease dose (15 U mL^-1^) has been reported to effectively degrade different forms of extracellular nucleic acids including G-quadruplexes^32^.

### Extraction and quantification of extracellular nucleic acids

Extracellular nucleic acids (eNAs) were isolated using a low-speed centrifugation protocol adapted from Nagler et al. (2018)^36^. Cultures were vortexed briefly, and centrifuged (10 min, 5000 × g) and the supernatants were filtered (0.22 µm) and combined with half a volume of sodium acetate buffer (3 M, pH 5.2). The cell pellets were washed with sodium acetate buffer containing 5 mM EDTA, and re-centrifuged, and the supernatants were pooled with the extracellular fractions. Nucleic acids were precipitated with double volume of 96% ethanol at -20°C overnight. The nucleic acids were pelleted by centrifugation (15 min 14000 x g), washed with 70% ethanol, air dried, and resuspended in TE buffer. eNAs were quantified fluorometrically with Qubit RNA, dsDNA, and ssDNA assay kits (Invitrogen) and spectrophotometrically using a Nanodrop instrument.

### Fluorescence staining and immunolabelling of extracellular nucleic acids

Extracellular nucleic acids in *M. barkeri* MS were visualized using fluorescence stains and nucleic-acid-specific antibodies. Live cultures were stained with TOTO-1 (1 µM), FM4-64 (10 µg mL^-1^), and SYTO-60 (20 µM). For immunolabeling of G4 quadruplexes, samples were incubated for 60 min in phosphate-buffered saline (PBS) containing 3% bovine serum albumin (BSA) with Atto488-BG4 anti-G4 antibody (1:100 dilution; Absolute Antibody, clone BG4, product code Ab00174-24.1). To detect B-DNA, samples were incubated for 60 min with an unconjugated mouse anti-B-DNA primary antibody (1:100 dilution in 3% BSA in PBS; VWR, clone AE-2, product code ANTIA250938-100) followed by Alexa Fluor 405 goat anti-mouse IgG (H+L) cross-adsorbed secondary antibody (AB2, 1:100 dilution in 3% BSA in PBS; Thermo Fisher, product code A-31553). Samples were counterstained with the membrane dye, FM4-64 (10 µg mL^-1^) and imaged using confocal laser scanning microscopy (CLSM, Zeiss LSM700) with 10x and 63x/1.4x objectives as described by Ajunwa et al. (2024)^26^. Abiotic controls were run alongside. For cells grown on cathodes, carbon-felt electrode samples were processed directly using the same immunolabeling protocol.

### *In vitro* DNAzyme/RNAzyme activity assays

DNAzyme/RNAzyme activity was assessed via peroxidation of ABTS (2,2’-azino-bis-(3-ethylbenzothiazoline-6-sulfonic acid)) using UV-VIS spectroscopy. Oxidized ABTS was generated *in vitro* by G4-hemin acting as an RNAzyme/DNAzyme and catalyzing a peroxidation reaction. Native extracellular G4-eNAs, synthetic G4-RNA, and G4-DNA (100 nM each) were incubated with hemin (100 nM) in MES buffer (25 mM MES, 200 mM NaCl, 10 mM KCl, 1% DMSO, 0.05% Triton X-100, pH 5.1) at 25°C for 30 min^45^. Reactions were initiated by adding ABTS (2 mM) and H_2_O_2_ (2 mM). Absorbance spectra (400-800 nm) of oxidized ABTS were recorded after 2 min using quartz cuvettes.

### Visualization of extracellular DNAzyme/RNAzyme activity *in vivo*

G-quadruplexes associate with hemin (iron-III-protoporphyrin IX) to form peroxidase-mimicking DNAzymes/RNAzymes^26,45^. Their activity catalyzes the transfer of electrons for the reduction of hydrogen peroxide (H_2_O_2_), which is read out by tyramide signal amplification (TSA). In this study, DNAzyme/RNAzyme activity was used to locate native extracellular G4s on the cell surface of *M. barkeri* and verify functional folding. *M. barkeri* culture was incubated with hemin (5 µM) in modified MES (2-(N-morpholino)-ethanesulfonic acid) buffer (pH 6.5), for 30 min at room temperature with gentle shaking (50 rpm). Cells were pelleted and incubated with TSA mix, containing tyramide conjugated with Alexa Fluor 488 (1:100; Invitrogen, B40953), ATP (2 mM; Thermo Scientific, R0441), and H_2_O_2_ (0.1%; Sigma, 216763) in MES buffer, for 90 min at room temperature, with gentle shaking (50 rpm). Hemin-free samples served as negative controls. Samples were washed in MES buffer and counterstained with SYTO™ 60 (20 µM) and visualized using CLSM with simultaneous dual-channel acquisition as described by Ajunwa et al. (2024)^26^.

### Nuclease and synthetic G4-RNAeffects under non-EET conditions

Three commercial nucleases, i.e., DNase I (EN0521, ThermoFischer) (500 U mL^-1^), RNase A (EN0531, ThermoFischer) (50 µg mL^-1^), and micrococcal nuclease (EN0181, ThermoFischer) (15 U mL^-1^)^32^, were tested for the effects on *M. barkeri* growth under non-EET conditions (methanol-grown). CH_4_ production and optical density (OD_600_) of the culture were monitored periodically and compared with untreated controls.

Separately, prefolded synthetic G4-RNA oligonucleotide (2.5 µM) was added to methanol-grown cultures to assess its impact on growth and methanogenesis under non-electrochemical conditions. G4-RNA was pre-folded by heating in GQ buffer (10 mM Tris and 100 mM KCl, pH 7.5) at 70°C for 5 minutes, followed by incubation on ice for 2 hours, and stored at -80°C^45^. CH_4_ production and optical density (OD_600_) of the culture were recorded periodically and compared with control without G4-RNA.

### Cell adhesion assays

To evaluate the contribution of eNAs to physical adhesion, *M. barkeri* was cultivated on carbon felt under anaerobic conditions. During active methanogenesis, micrococcal nuclease (15 U mL^-1^) or buffer control was added and further incubated for an additional 12 h. Following treatment, carbon felt was gently removed, briefly rinsed in PBS (pH 7.4) to eliminate loosely bound cells, and transferred into fresh sterile medium and monitored for methane production.

For quantitative assessment of adhesion, biomass retained on the carbon felt was measured as total protein. Adhered cell-protein was extracted by immersing the felt in 2% CHAPS in PBS and boiling for 5 min. Lysates were analyzed by a bicinchoninic acid (BCA) assay (Thermo Scientific, Cat. No. 23225) according to the manufacturer’s protocol. Total protein concentrations served as proxy for adhered biomass, while methane production by felt-bound cells ± micrococcal nuclease served as a functional readout of nuclease impact.

### Electrochemical Impedance Spectroscopy (EIS) analysis of *M. barkeri*

*M. barkeri* cells were harvested anaerobically, washed twice with fresh media and the concentrated (400 µL) cells were transferred to modified Screen Printed Electrode (SPE) reactors for EIS analysis. For SPEs, potentials were converted to SHE based on calibration against known redox behavior under identical experimental conditions. Reactors were incubated at 37°C under static conditions for 30 min prior to measurements to allow stabilization of the cell-electrode interface. Pre-folded artificial G4-RNA was added to the reactors at a final concentration of 5 µM. Abiotic controls, abiotic samples with added G4-RNA, cell-only samples, and cells supplemented with G4-RNA were prepared in four independent replicates each (n=4).

Cyclic voltammetry (-0.6V to 0.4V v SHE) was employed to characterize the redox activity and electrochemical reversibility of *M. barkeri*-modified electrodes to check both the nature of redox potential for electron transfer and the stability of the cell-electrode interface under operating conditions. The CV, chronoamperometry and electrochemical impedance spectroscopy (EIS) were performed in two sequential runs. Chronoamperometry was conducted at an applied potential of -430 mV vs SHE for a duration of 6 h, followed by EIS measurements. EIS measurements were conducted after an equilibration period of 60s using a fixed DC bias. The applied DC potential (E_dc) was -430 mV vs SHE with a sinusoidal AC perturbation amplitude (E_ac) of 5 mV. Impedance spectra were recorded over a frequency range from 100 kHz to 0.1 Hz using a frequency scan mode. Data were fitted using MultiTrace4.5, PalmSens, and the fitted parameter data are presented as mean ± s.d. of four independent replicates. Impedance spectra were confirmed as Kramers-Kronig consistent.

### Identification of putative G4 sequences in the genome of *M. barkeri*

Putative G4-forming sequences in the *M. barkeri* MS genome were identified using the G4Hunter algorithm^46^. The analysis parameters included a window size of 25 and a threshold score of 1.2, ensuring the accurate detection of potential G4 motifs^46^. The frequency of G4 motifs was quantified across the whole genome, coding sequences, promoters, tRNA, and rRNA regions. Promoter regions were defined as the 2000 bp sequences upstream of coding regions. G4 motif density was calculated by dividing the total number of sequences by the genome length.

## Acknowledgements

This is a contribution to a Novo Nordisk Foundation Ascending Investigator grant NNF21OC0067353 to AER. We would like to express our gratitude to the Nordcee technicians, particularly Lasse Ørum Schmidt and Anni Glud for laboratory assistance. We are also thankful to Bo Thamdrup, Don Canfield, Ronnie Glud for their comments on an earlier summary of our work and all members of the Rotaru lab for helpful discussions.

## Author contributions

Conceptualization: AER, RK, RLM

Methodology: RK

Investigation: RK, OMA, SK

Visualization: RK, AER

Funding acquisition: AER, RLM

Project administration: AER

Supervision: AER, RLM

Writing – original draft: RK, AER

Writing – review & editing: RK, OMA, SK, RLM, AER, HB, JTB

## Data and materials availability

All data needed to evaluate the conclusions in the paper are present in the main text and/or the supplementary materials. Source numerical data underlying all figures and tables are available from the corresponding authors upon reasonable request.

